# Chromatin opening ability of pioneer factor Pax7 depends on unique isoform and C-terminal domain

**DOI:** 10.1101/2023.03.01.530660

**Authors:** Virginie Bascunana, Audrey Pelletier, Arthur Gouhier, Amandine Bemmo, Aurelio Balsalobre, Jacques Drouin

## Abstract

Pioneer factors are transcription factors (TFs) that have the unique ability to recognise their target DNA sequences within closed chromatin. Whereas their interactions with cognate DNA is similar to other TFs, their ability to interact with chromatin remains poorly understood. Having previously defined the modalities of DNA interactions for the pioneer factor Pax7, we have now used natural isoforms of this pioneer as well as deletion and replacement mutants to investigate the Pax7 structural requirements for chromatin interaction and opening. We show that the GL+ natural isoform of Pax7 that has two extra amino acids within the DNA binding paired domain is unable to activate the melanotrope transcriptome and to fully activate a large subset of melanotrope-specific enhancers targeted for Pax7 pioneer action. This enhancer subset remains in the primed state rather than being fully activated, despite the GL+ isoform having similar intrinsic transcriptional activity as the GL-isoform. C-terminal deletions of Pax7 lead to the same loss of pioneer ability, with similar reduced recruitments of the cooperating TF Tpit and of the co-regulators Ash2 and BRG1. This suggests complex interrelations between the DNA binding and C-terminal domains of Pax7 that are crucial for its chromatin opening pioneer ability.

## Introduction

Pioneer transcription factors are unique in their ability to recognise and bind their target DNA sequences in closed chromatin (1). Their action initiates chromatin opening by a process that has received recent insight (2). This unique property has endowed this class of transcription factors with the power to direct new cell fates in development (3,4). This is achieved by mostly targeting new enhancer repertoires for chromatin opening and activation of novel transcriptomes in daughter compared to mother cells (5-8). Hence, the activation of new transcriptomes defines new cell identities. Pioneers also establish an epigenetic memory of these new cell identities through demethylation of target enhancer DNA (1,8-11). It appears as if pioneer ability may not be required beyond the initial chromatin opening and establishment of epigenetic memory: beyond this epigenetically stable process, pioneer factors often serve as regular transcription factors to maintain differential cell specific transcriptomes (12). In principle, pioneer factors may elicit these separate functions through unique and different protein domains and co-factors (13). The identification of unique features within pioneer factors that account for their pioneering compared to transcriptional functions would in itself serve to better define the unique aspects of chromatin opening by pioneer factors.

The structure of a few pioneer factors was probed through structure function studies. For the FOXA factors that specify endoderm derivatives, closed chromatin binding was shown to involve the DNA binding domain (DBD) that interacts with its target DNA sequences, as well as the C-terminal domain that interacts with histones H3 and H4 (14). Further, the FoxA DBD is flanked by wings that have structural similarities to histone H1 and that may contribute to chromatin interactions by mimicry/displacement of histone H1 (15). The C-terminal domain of the pioneer EBF1 involved in B cell identity was shown to be essential for closed chromatin interaction and pioneering (7). More recently, an intrinsically disordered region of the pioneer PU.1 was shown to be important for interaction with linker histone H1-compacted nucleosome arrays (16).

The pioneer factor Pax7 is expressed in a variety of tissues, but its pioneering role is clearly supported only in the pituitary gland where Pax7 opens a new enhancer repertoire to specify the intermediate pituitary tissue and the melanotrope cell fate (8,17). This activity requires the co-recruitment and cooperation with a nonpioneer factor, the Tbox factor Tpit (18). Elsewhere, Pax7 is expressed in skeletal muscle and specifies subdomains of the neural tube (19,20). In the myogenic lineage, the related Pax3 and Pax7 transcription factors are critical to implement the myogenic fate and in particular for the transition of progenitors into differentiated myogenic cells (21). A clear indication of Pax7 pioneering in these latter systems remains poorly established (21,22), because demonstration of the pioneering ability requires access to the tissues or cells of origin prior to chromatin opening by the pioneer factor: this is not always easily accessible as an experimental system.

In the present work, we have investigated the structural features of Pax7 implicated in chromatin opening of the enhancer repertoire that specifies the pituitary melanotrope fate. The ability to reprogram a corticotrope cell model, the AtT-20 cells, into melanotropes enabled us to investigate the pioneer ability of naturally occurring isoforms of Pax7, together with a serie of deletion and alanine replacement mutants. The Pax7 isoforms differ by just a few amino acids within the DNA binding paired domain but these suffice to create isoforms with or without pioneer ability, but with similar inherent transcriptional activities. The pioneering-deficient Pax7 isoform is similar to the related Pax3 in their inability to activate the melanotrope transcriptome. A similar loss of pioneering ability was observed for C-terminal deletion mutants of Pax7. It is striking that similar losses of pioneering ability are produced by either introduction of two amino acids within the paired domain or by deletion of C-terminal sequences. The loss of pioneering ability is clearly linked to the loss of chromatin opening rather than to failure of chromatin recruitment. This pattern corresponds to an inability of one isoform and mutants to proceed to the second step of chromatin opening by Pax7 (2); hence, these forms can only prime but not fully activate a subset of target enhancers that are associated with the melanotrope transcriptome.

## Materials and Methods Cell culture

AtT-20/D16v-F2 cells were cultured in Dulbecco’s modified Eagle’s medium supplemented with 10% fetal bovine serum and antibiotics (penicillin/streptomycin). To generate stable transgenic AtT-20 Pax7 cell populations, retroviruses were packaged using the Platinium-E Retroviral Packaging Cell Line (Cell Biolabs, catalog #RV-101) and infections performed as described (23). Selection of retrovirus-infected cell populations was achieved with 400 g/mL Geneticin (Gibco, #11811-031). Resistant colonies were pooled to generate populations of hundreds of independent colonies.

### Luciferase assays

AtT-20 cells were plated in 12-well dishes (4 × 10^5^ cells per well) and transfections performed the next day using PEI 25K (Polysciences #23966-1) in triplicates. Transfected DNA (total 2 μg in 150 μL DMEM) was added to the PEI 25K solution (4 μL per 150 μL DMEM) and incubated at room temperature for 30 minutes before adding 100 μL of the mixture to each well. The media was changed 24 hours after transfection and luciferase activity assessed 48 hours after transfection. Growth medium was removed from transfected cells and 200 μL of lysis buffer (Tris 100mM, NP40 0.5% and DTT 0,001M) added to each well. After 10 minutes on a vigorous shaker, 100 μL of cell lysate supernatant was used for analysis using the GloMax Navigator luminometer (Promega) using the kinetics protocol (100 μL injection, speed 100 μL/s, integration time 10 s). Statistical significance was assessed by 2-way ANOVA and Tukey’s multiple combinations test.

### ChIP-Seq, ATAC-Seq

ChIP experiments were performed in AtT-20 cells as described earlier (8,24). ATAC-Seq was performed as described in (8). The DNA libraries and sequencing flow cells were prepared by the IRCM Molecular Biology Core Facility following the recommendations of Illumina (Illumina, San Diego, California). ChIPs samples were sequenced on the Illumina Hi-Seq 2000 sequencer. Supplementary Table 1 provides details on the antibodies used for ChIPs. ChIP-Seq and ATAC-Seq average profiles were generated by the average signal function of Easeq software (http://easeq.net) and the data exported to Microsoft Excel to create graphs. To view the ChIP-Seq and ATAC-Seq heatmaps, we used the Integrative Genome Viewer tool of Easeq (25) or a custom Python program (13). All ChIP-Seq and ATAC-Seq data except Flag ChIPs were normalised relative to Constitutive and random sites.

### Nuclear extracts

To prepare nuclear extracts, AtT-20 cells from a 100 mm Petri dish (15-20 ×10^6^ cells) were washed and collected in PBS. The cells were resuspended in 800 μL of a cold buffer with low salt content (10 mM HEPES pH 7.9, 10 mM KCl, 0.1 mM EDTA, 0.1 mM EGTA, 1 mM DTT, 0.5 mM PMSF, 0.6 μM aprotinin, 0.6 μM leupeptin and 2 μM pepstatin A). After 15 minutes of incubation on ice, 100 μL NP40 10% was added. Cytosolic fractions were removed, and nuclear proteins extracted in 100 μL of high salt cold buffer containing 20 mM HEPES pH 7.9, 400 mM KCl, 1 mM EDTA, 1 mM EGTA, 1 mM DTT, 1 mM PMSF, 0.6 μM aprotinin, 0.6 μM leupeptin and 2 μM pepstatin A. Protein concentration was quantified in the Bradford assay.

### EMSA

Double-stranded DNA probes were described and labeled as in Pelletier et al (13). Gel shift assays were performed using 1 μg of nuclear protein extracts from different AtT-20 cell lines pre-incubated 10 min on ice with 1 μg of nonspecific carrier DNA poly(dI-dC) and poly(dA-dT) in binding buffer (25 mM HEPES pH 7.9, 84 mM KCl, 10% glycerol, 5 mM DTT). For purified polypeptides, 300 fmoles were used per binding reactions. Equal amounts of radiolabeled DNA probes (50,000 cpm) were then added for a total volume of 20μL and incubated 60 min on ice. If applicable, 1 μg of Flag M2 antibody (F3165, SIGMA) was added to the mixture during the last 10 minutes of incubation.

Nondenaturing polyacrylamide gels (40 mM Tris, 195 mM glycine, 0.08% APS, 0.5 μL/mL TEMED, 5% Acrylamide:Bisacrylamide in 19:1 ratio), were prerun in Tris-glycine buffer (40mM Tris and 195 mM glycine) for 60 min at 300 V at 4°C. The gel shift reaction mixture was then loaded, and the gel run for 3 h at 300 V at 4°C. Gels were dryed on 3MM Whatman paper in a gel dryer at 80°C during 60 min under vacuum. Autoradiography films (HyBlot CL, catalog # E3018) were then exposed with the dried gels for 10 to 48 h using an intensifier screen.

### RT-qPCR

RNA was extracted from AtT-20 cells using the RNeasy Mini kit (Qiagen #74104), and cDNA synthesized using 5 μg RNA and SuperScript III reverse transcriptase (Invitrogen, #18080-044) following manufacturer’s recommendations. The resulting cDNAs (5 μl of 1/50 dilution) were analyzed by qPCR using SYBR Green reagent (ThermoFisher Scientific #A25741) supplemented with 500 nM of each gene-specific primer pair (provided in Supplementary Table 2) in a total volume of 10 μl. At least 2 biological replicates were analyzed in duplicates for each condition using a ViiA™ 7 Real-Time PCR device (ThermoFisher Scientific), and results were analyzed using the accompanying software. Gene expression values were normalized relative to the TBP transcript and statistical significance assessed by 2-way ANOVA and Tukey’s multiple combinations test.

### ChIP-qPCR

ChIP experiments were performed in AtT-20 cells as described (8). Immunoprecipitated DNAs were diluted in 100 uL and were analyzed by qPCR using the SYBRgreen reagent (ThermoFisher Scientific #A25741) completed by 500 nM of each enhancer-specific primer pair (provided in Supplementary Table 3) in a total volume of 10 μl. At least 2 biological replicates were assessed in duplicates for each condition using a real-time ViiA™ 7 (ThermoFisher scientific) PCR device, and the results were analyzed using the associated software. ChIP recruitment values were normalized to those of two random loci (H3K9me3poor, H3K9me3rich as listed in Supplementary Table 3).

### NGS data analysis

ChIP-seq and ATAC-seq paired-end sequenced reads were trimmed using Trimmomatic/0.36

(26) and aligned to the mouse mm10 reference genome using Bowtie/2.3.5 (--very-sensitive --no-mixed --no-unal) (27). Bam files were created using view from SAMtools/1.9 (28) and duplicated reads were removed using MarkDuplicates from Picard/2.17.3 (Broad Institute). Coverage tracks were created using bamCoverage (--normalizeUsing RPKM -bs 10 -e) from deepTools/3.3.0 (29). Peaks were called against sequenced input DNA or control Flag ChIP using callpeaks (-f BAMPE -p (1e-3) -g mm) from MACS/2.1.1 (30). Pax7 recruitment site categories are based on previously published lists (13). In brief, Pax7 ChIP-seq peaks were categorized as “Pioneered” if they had no signal before, and gains (p-value < 1e-5 for ChIP and < 1e-3 for ATAC) of H3K4me1, p300 and ATAC-seq, after Pax7 expression. “Activated” sites gained p300 after Pax7 and Constitutive sites where already positive for all marks before Pax7. To determine H3K4me1 peak sub-categories, H3K4me1 signals were quantitated within a 200pb window by easeq around the summit of the Pax7 peaks. We next compared these with regions at 500bp on either side of the Pax7 summits to determine whether they are equal to, lower than, or greater than, the central region of the Pax7 peaks.

### Motifs analysis

The motifs recognized by Pax7 were described by Pelletier et al (13). Presence or absence of a specific Pax7 motif within 50 base pairs on each side of the Pax7 ChIP summit at pioneer sites was assessed using the command Homer findMotifsGenome.pl (http://homer.ucsd.edu/homer/ngs/peakMotifs.html). The number of Pax7 motifs present within a window of 200 base pairs around the summit of each pioneer site was assessed using a custom program based on Python (13).

## Results

The pioneer factor Pax7 is expressed as different isoforms that result from differential exon splicing (20,31). Prior work defined the nature of isoforms expressed in muscle (19,31) and brain (32,33) tissue and we assessed the relative expression of Pax7 isoforms in the pituitary gland where we previously characterized the pioneer ability of Pax7 (8,17,18). The Pax7 isoforms are characterized by the differential insertion of either a single or a pair of amino acids at two positions within the DNA binding paired (PD) domain. The first differential splicing involves a glutamine (Q) residue inserted before helix 4 of the helix-turn-helix (HTH) RED domain (Q+) whereas the second differential splicing event involves a glycine-leucine pair inserted (GL+) at the beginning of helix 6 of the same RED HTH domain (Figure 1a). In the pituitary intermediate lobe, the predominant isoform as assessed by RT-qPCR is the Q+GL-isoform that represents about 75% of the mRNA whereas the remainder is the Q+GL+ isoform, with no expression of the Q-isoforms (Figure 1a). The Pax7 Q+GL-isoform was used in our prior work (8,13,17,18) and constitutes the reference isoform in the present work.

**Figure 1.**
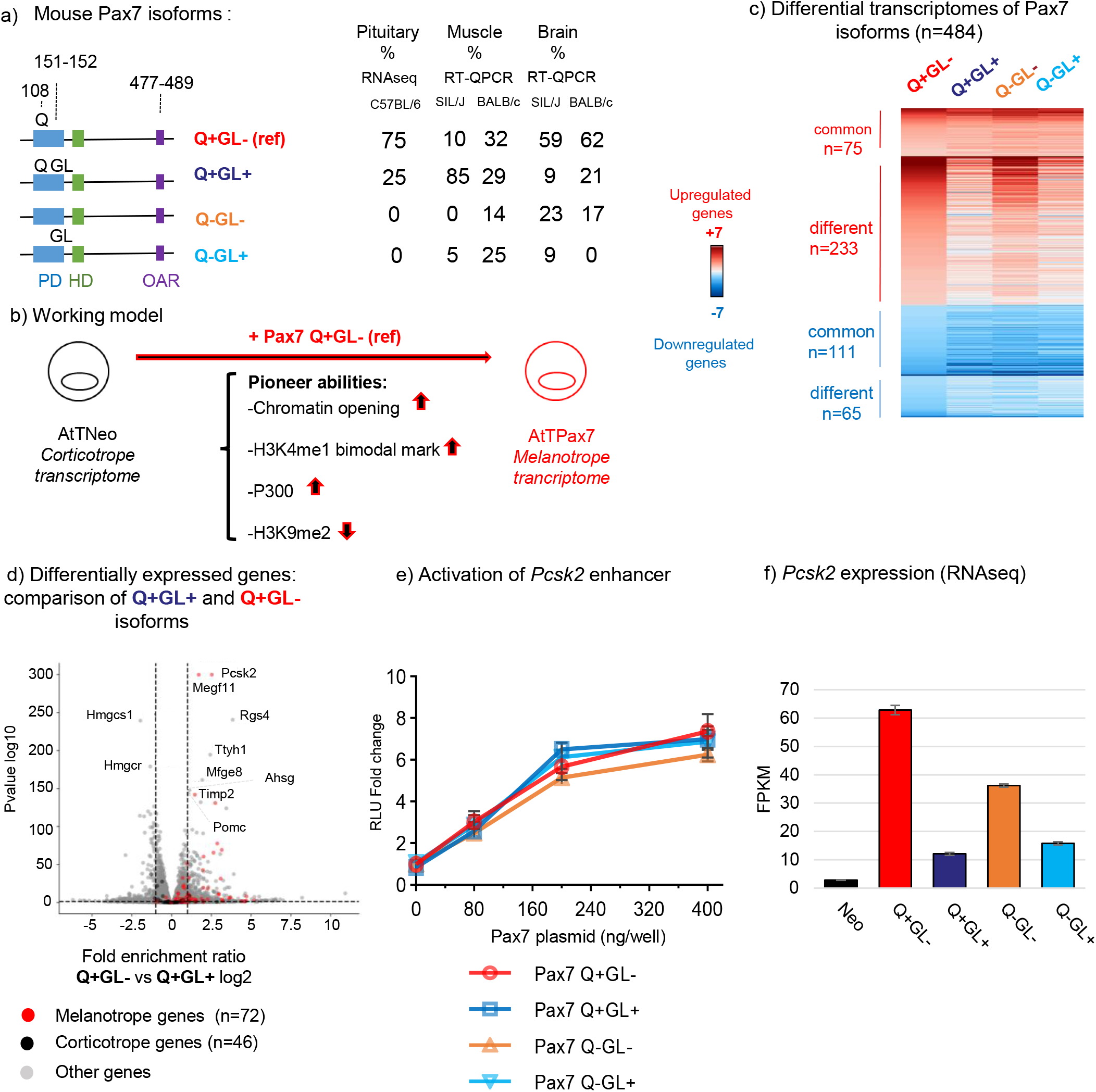
Isoform-specific activation of the Pax7-dependent melanotrope transcriptome. **a**) Differential splicing produces naturally occurring isoforms of mouse Pax7 with a glutamine (Q) and/or glycine and leucine (GL) inserted within the DNA-binding paired domain (PD, blue). The reference Pax7 isoform (Q+GL-) used in the present and in previous work (8,13,17,18) is the predominant isoform expressed in the mouse pituitary. The relative abundance of mRNAs for different isoforms was measured in tissues of different mouse strains by either RNA-Seq or RT-qPCR as indicated. Data from mouse muscle and brain (31) were previously published. **b**) The mouse AtT-20 cells are used as a model of pituitary corticotropes and expression of Pax7 reprograms these towards the melanotrope fate (17) by opening chromatin at a repertoire of melanotrope enhancers (8,17). **c**) Differential transcriptomes of Pax7 isoforms expressed in AtT-20 cells was determined by RNA-Seq. The heatmaps represent differentially expressed genes compared to control AtTneo cells for each of the four Pax7 isoforms. Activated and repressed genes are indicated in red and blue, respectively. **d**) Volcano plot comparison of differentially expressed genes in AtT-20 cells expressing the Q+GL+ isoform compared to the reference Q+GL-isoform. Highlighted melanotrope (red dots) and corticotrope (black dots) genes reflects the ability of the reference Pax7 isoform to activate the melanotrope transcriptome and to repress genes of the corticotrope transcriptome as previously defined (8). **e**) Transcriptional activity of different Pax7 isoforms was assessed using a luciferase reporter driven by the melanotrope-specific *Pcsk2* gene enhancer (17). **f**) *Pcsk2* gene expression assessed by RNA-Seq analysis of AtT-20 cells expressing each Pax7 isoform as indicated.

### GL+ isoforms fail to activate the melanotrope transcriptome

In order to assess the properties of the Pax7 isoforms, we used the gain-of-function system previously developed (Figure 1b) in mouse AtT-20 cells that represent a model of pituitary corticotrope cells and that undergo reprogramming into melanotropes following Pax7 expression (17). Pools of AtT-20 cells each expressing a different Pax7 isoform under control of retroviral sequences were obtained and shown to express similar levels of PAX7 proteins (Supplementary Figure 1a). Complete transcriptomes were obtained for these cells by RNA-Seq and analyzed for differentially expressed genes (Figure 1c and Supplementary Figure 1b). Comparison of the transcriptomes reveals that GL+ isoforms affect expression of far fewer genes than the GL-isoforms; further, the presence of Q (Q+ isoforms) has a marginal effect (Figure 1c). Significantly, volcano plot representation of differentially expressed genes between the Q+GL- and Q+GL+ isoforms reveals that most (91%) previously identified (8) melanotrope genes are preferentially activated in Q+GL-compared to Q+GL+ cells (Figure 1d and Supplementary Figure 1c,d). The ability of Pax7 to implement this melanotrope program of gene expression thus appear to be mostly dependent on the GL-isoforms and to be prevented by the insertion of GL in the PD domain of the GL+ isoform.

This differential gene expression could be due to differences in transcriptional activation capability, and this was assessed directly by co-transfection of the various isoforms with reporter plasmids containing either the natural enhancer of the *Pcsk2* gene (Figure 1e) or reporters containing multimers of various Pax7 DNA binding sites (13) (Supplementary Figure 1e). The different Pax7 isoforms have very similar transcriptional ability in all these systems. Hence, it is not transcriptional ability that accounts for the unique properties of the isoforms. In striking contrast, the levels of the melanotrope-specific *Pcsk2* mRNA are very different in cells expressing different isoforms (Figure 1f). Indeed, the GL-isoforms express significantly higher levels of *Pcsk2* mRNA than the GL+ isoforms with the greatest difference being between the two isoforms expressed in pituitary, namely the predominant Q+GL-compared to the Q+GL+ isoform. Expression of *Pcsk2* was previously shown to depend on the Pax7 pioneer ability (8,17).

### All Pax7 isoforms exhibit similar genomic occupancy

Since the intrinsic transcriptional activity of the Pax7 isoforms does not explain their differential effect on mRNA expression, we assessed the ability of each isoform to be recruited to chromatin and trigger chromatin opening. We first performed ChIP-Seq for Pax7 recruitment in cells expressing the different isoforms and globally all isoforms showed similar binding (Figure 2a and Supplementary Figure 2). Subtle differences appeared however when Pax7 recruitment is compared at previously described subsets of enhancers (8) (Figure 2b), namely Constitutive (open and active before Pax7), Activated (gain of ATAC, H3K4me1 and p300 signals after Pax7 binding with prior weak signals) and in particular at Pioneer enhancers (that had no active marks before Pax7 but gained marks after). Whereas binding was similar for all isoforms at Constitutive and Activated enhancers, recruitment of the Q+GL+ isoforms is impaired, but not lost, at Pioneer enhancers compared to the reference Q+GL-isoform (Figs. 2c and 2d), except for a small group of 115 Pioneer sites (13%, labelled as GL-sensitive binding) where binding is lost (Figure 2c).

**Figure 2.**
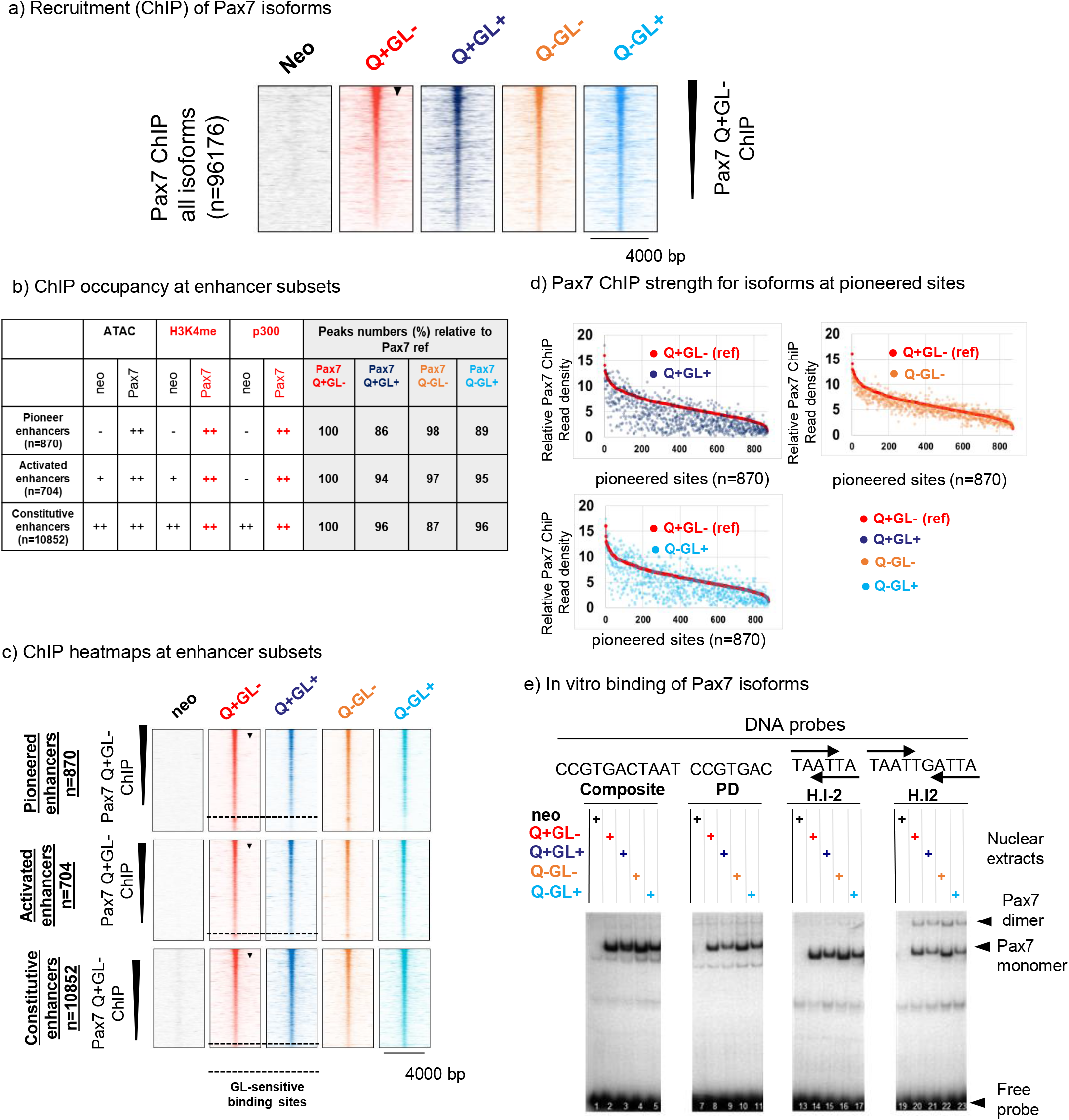
Similar genomic recruitment of all Pax7 isoforms. **a**) ChIP-Seq analysis of genomic recruitment for Pax7 isoforms as indicated. Heatmaps are ordered by decreasing intensity for the reference Q+GL-isoform (indicated by arrowhead). **b)** Pax7 isoform occupancy (in comparison to reference isoform) at the different subsets of Pax7 recruitment sites as indicated. The presence/absence of different marks used to define enhancers are indicated for sites pioneered or activated by Pax7 in comparison to Constitutive enhancers that are not affected by Pax7 occupancy as defined in ref.(13). **c)** Heatmap representation of Pax7 isoform recruitment at the indicated subsets of Pax7 recruitment sites rank-ordered relative to the reference Q+GL-isoform (arrowhead). A small subset of about 13% of Pioneer sites no longer binds the GL+ isoforms as indicated by the dashed line at bottom of heatmaps. **d**) Comparisons of Pax7 ChIP recruitement strength for the indicated isoforms at the subset of Pioneer sites. **e**) In vitro binding of Pax7 isoforms to various DNA target sites (probes) for either Composite, paired (PD), or HD (H.I-2, H.I2) domains was assessed by electrophoretic mobility shift assay (EMSA) as described previously (13). The positions of Pax7 monomers and dimers, as well as free probe, are indicated on the right.

The small decrease in genomic recruitment of the GL+ isoforms could be due to intrinsic DNA binding ability, and this was assessed by *in vitro* binding in electrophoretic mobility shift assay (EMSA) experiments (Figure 2e). We previously (13) characterized Pax7 binding to various DNA target sites for either or both of its DNA binding paired (PD) and/or homeo (HD) domains (Figure 2e). Hence, we assessed binding of the Pax7 isoforms to these probes and the experiments showed similar *in vitro* binding ability for the various isoforms with a slightly weaker binding of the GL+ compared to the GL-isoforms: this was true for all probes studied (Figure 2e). It is thus possible that the weaker recruitment observed for the GL+ isoforms in ChIP experiments may in part be due to lower intrinsic DNA binding affinity. However, the decrease in Q+GL+ recruitment compared to the Q+GL-isoforms (Figure 2c) appears greater than expected from differences in *in vitro* binding.

### Melanotrope enhancers are primed rather than activated by GL+ isoforms

In order to assess the ability of Pax7 isoforms to initiate chromatin opening, we performed ATAC-Seq on cells harboring the pituitary-expressed Q+GL- and Q+GL+ isoforms. Whereas similar ATAC-Seq signals are observed at Constitutive sites for both isoforms, the Q+GL+ isoform is unable to open a subset of sites in both the Pioneer and Activated subcategories (Supplementary Figure 3 a-e). It is ∼45% of the Pioneer sites that fail to show significant ATAC-Seq signal in presence of Q+GL+ compared to Q+GL-isoform. We assessed whether the underlying Pax7 DNA binding motifs present in each subset of enhancers could account for the sensitivity to insertion of the GL residues in the PD domain. Neither frequency of individual motifs (Supplementary Figure 3f) nor total number of Pax7 binding motifs (Supplementary Figure 3g) can explain the difference between the properties of the two Pioneered subsets.

To further assess Pax7-dependent deposition of active enhancer marks, we performed ChIP-Seq for H3K4me1, a mark of potentially active enhancers. Interestingly, the subset of enhancers that did not show significant ATAC-Seq signals, do show a small gain in H3K4me1 that mostly presents as a single weak peak compared to the bimodal H3K4me1 distribution typically observed at fully active enhancers (Supplementary Figure 3c). This single peak H3K4me1 profile is typical of Primed enhancers and hence the subset of enhancers that are impaired by the presence of GL+ fails to be fully activated but is nonetheless remodelled from the inactive to the Primed state. Since this change in H3K4me1 profile is the most significant GL-dependent alteration, we quantitated the H3K4me1 profiles as single or bimodal profiles, the latter subgroup being further subdivided as symmetric or asymmetric bimodal profiles (Figure 3a-b). Examples of these profile changes are shown for two Pax7-dependent enhancers (Figure 3d). The H3K4me1 profiles thus provide the best illustration of the impact of GL insertion for pioneer action: whereas a subset of sites is insensitive (GL-insensitive), three subsets that retain Pax7 binding with decreased ATAC signal (Figure 3c) exhibit a shift from bimodal to single peak (either symmetrical or asymmetric) H3K4me1, indicative of a failure of nucleosome displacement (Figure 3b,e).

**Figure 3.**
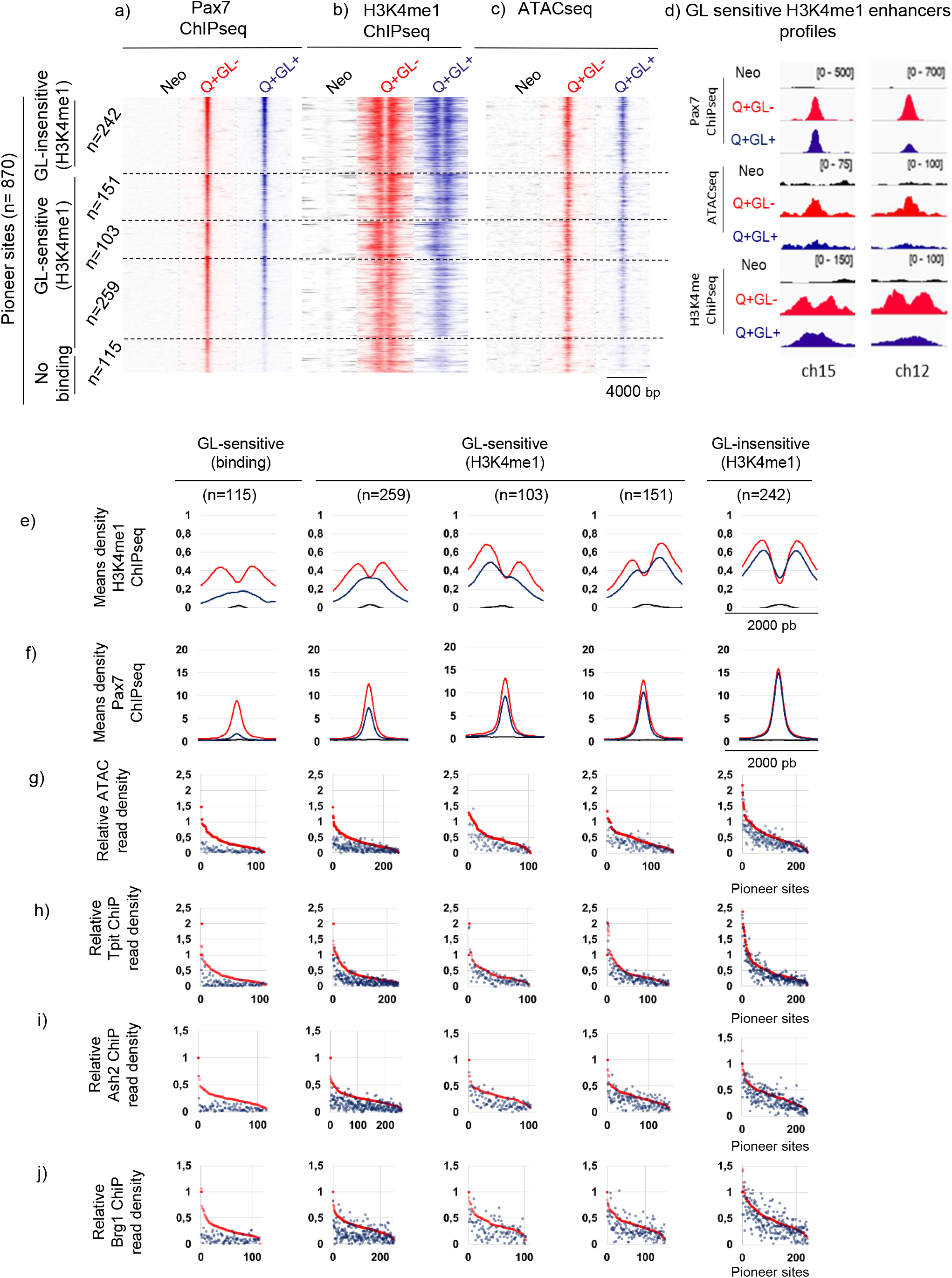
The melanotrope-specific Pax7 enhancers are primed rather than activated by the GL+ isoform. **a-c**) Pioneer ability of the two pituitary-expressed Pax7 isoforms (throughout, GL- and GL+ isoforms are shown in red and blue, respectively) assessed by Pax7 ChIP-Seq (a), H3K4me1 ChIP-Seq (b) and ATAC-Seq (c). Heatmaps are shown for the subsets of Pioneer enhancer sites that are subdivided into GL-sensitive, GL-insensitive and No binding subsets based on the H3K4me1 profiles and Pax7 recruitment. **d**) Gene browser views of Pax7 ChIP-Seq, ATAC-Seq and H3K4me1 ChIP-Seq signals for control AtT-20 cells (Neo) and cells expressing the indicated Pax7 isoform at two representative loci (chr15:62959658-62959772 and chr12:42716335-42716480). **e, f**) Average profiles (relative reads densities) for the indicated marks determined by ChIP-Seq at the subsets of Pax7 Pioneer sites. **g-j)** Data distribution profiles are ordered relative to Pax7 GL-signals for the indicated marks and Pioneer subsets (a).

We then assessed how stable Pax7 binding may differ between the subsets of pioneered enhancers (Figure 3f) and whether this may account for the difference in chromatin opening revealed by ATAC-Seq (Figure 3g). While Pax7 binding is more affected for the GL-sensitive subsets compared to the GL-insensitive subset, significant binding remained for most sites within the subsets of GL-sensitive enhancers (Figure 3f). However, the loss of chromatin opening ability revealed by ATAC-Seq is significantly different between the subsets (Figure 3g). The analysis of transcription factor DNA binding motifs did not provide clues to explain the differences between GL subsets (Supplementary Figure 4).

We further compared the status of the GL-sensitive and GL-insensitive enhancers by performing ChIP-Seq for Tpit, a Pax7 cooperating nonpioneer (18), and chromatin remodelling proteins associated with Pax7 pioneering (8). Whereas recruitment of the Pax7-cooperating transcription factor Tpit and of the remodelling complex proteins Ash2 and Brg1 are similar for Q+GL+ and Q+GL-at the GL-insensitive subset of enhancers, their recruitment is about half for the GL-sensitive subset (n=259) of enhancers that exhibit a single peak of H3K4me1 (Figure 3 h-j and Supplementary Figure 5). While these recruitment decreases are correlated with weaker chromatin opening (ATAC), they are more strikingly associated with a failure of nucleosome eviction (single peak rather than bimodal H3K4me1 profiles) and enhancers that appear stalled in the Primed state (Figure 3e). This could be due to inadequate recruitment of Tpit and/or of the SWI/SNF complex containing Brg1.

### Pax3 (GL-) fails to open the GL-sensitive melanotrope specific enhancers

The Pax transcription factor that is most similar to Pax7 is Pax3 and indeed, these two factors cooperate for implementation of the skeletal muscle program (34). Even though Pax3 is not expressed in the pituitary, it is interesting to compare the pioneering ability of those two factors in view of their co-expression in other tissues. Pax3 was thus introduced into AtT-20 cells and expressed at similar levels as Pax7 (Supplementary Figure 6a, b). Pax3 and Pax7 have very similar sequence in their DNA binding domains and are relatively conserved in their C-terminal regions (Figure 4a and Supplementary Figure 6a), but Pax3 does not have the OAR sequence present in the C-terminus of Pax7. Pax7 and Pax3 have similar *in vitro* binding ability when assessed by EMSA (Figure 4b). However, AtT-20 cells expressing Pax3 showed altered expression of far fewer genes compared to Pax7 Q+GL- and strikingly, a similar subset of genes that are unaffected by the GL+ isoform of Pax7 also fail to be upregulated by Pax3 that is itself Q+GL-(Figure 4c). Comparison of Pax3/7 and H3K4me1 ChIP-Seq together with their ATAC-Seq profiles show that Pax3 behaves similarly to the Q+GL+ Pax7 isoform despite Pax3 being GL-(Figure 4 d-f). Hence, Pax3 is similarly deficient in chromatin opening ability (Figure 4e) despite limited impact on ChIP-Seq recruitment at the GL-sensitive subset (Figure 4d) together with a H3K4me1 single peak profile typical of Primed enhancers (Figure 4f).

**Figure 4.**
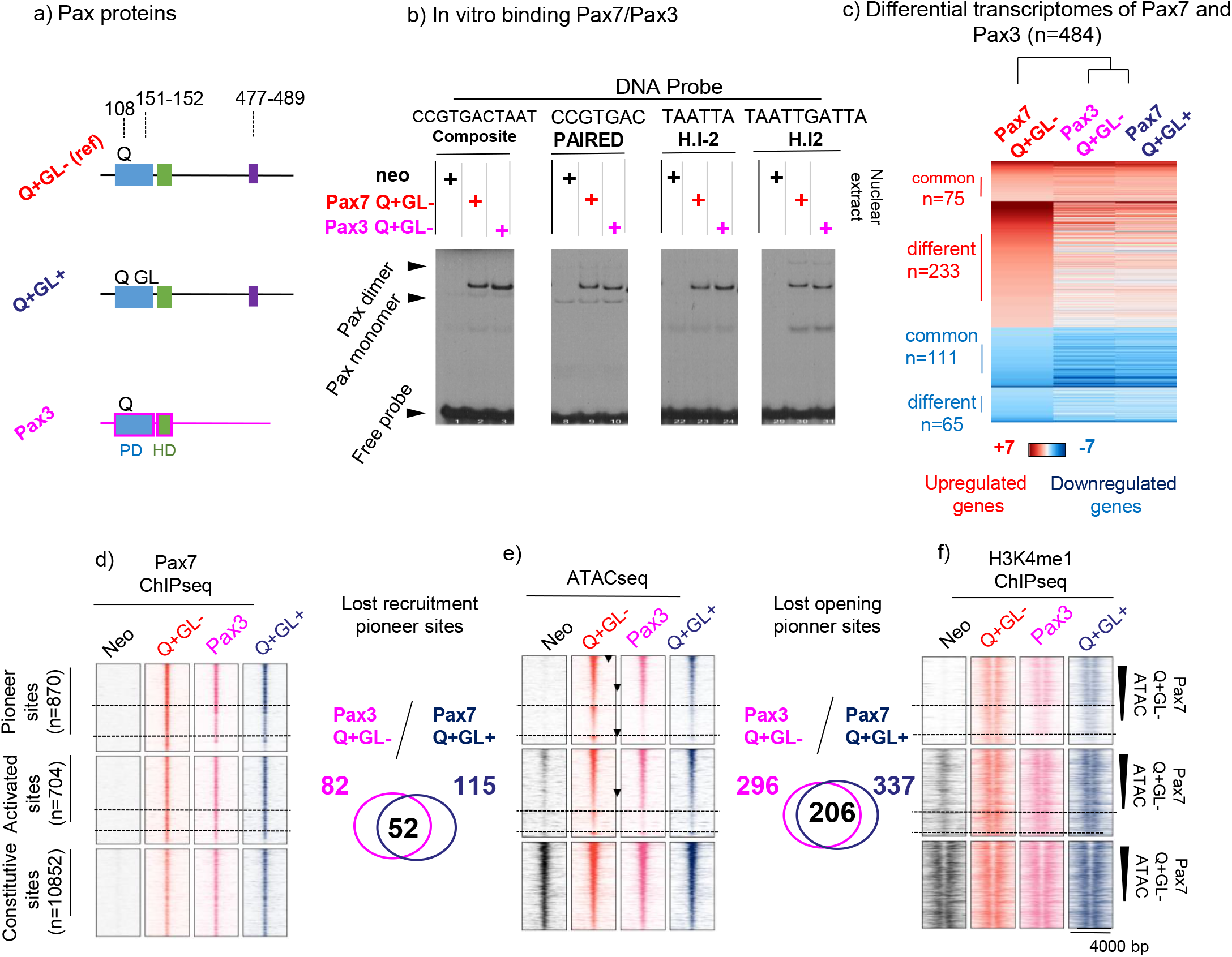
Pax3 has impaired pioneer activity comparable to that of Pax7 GL+. **a**) Schematic representation of Pax3 and Pax7 isoforms. Pax3 is Q+GL- and does not have the OAR-containing C-terminal extension present in Pax7. **b**) In vitro binding abilities (EMSA) of Pax3 and Pax7 using DNA probes for their different DNA binding sites. **c**) Differential transcriptome analyses of Pax3 and indicated Pax7 isoforms expressed in pituitary AtT-20 cells. Heatmaps represent differential transcriptomes compared to control AtT-20neo cells. **d-f**) Comparison of Pax3 action with Pax7 isoforms at different subsets of recruitment sites. Venn diagrams show significant overlap for Pax3 and Pax7 GL+ loss of sites compared to Pax7 GL-in ATAC-Seq (**d**) and in ChIP-Seq (**e**). Similar decreases are shown for H3K4me1 ChIP-Seq distributions (**f**).

### Pax7 C-terminus is required for recruitment to melanotrope specific enhancers

To define Pax7 domains involved in pioneer activity, we conducted C-terminal deletions of Pax7 (Figure 5a). The choice of C-terminal deletions endpoints was guided by alignment of the mouse Pax7 and Pax3 sequences in order to delete blocks of homology where possible (Supplementary Figure 6a). Pools of AtT-20 cells expressing these deletions mutants at similar levels (Supplementary Figure 6b) were first assessed for *in vitro* DNA binding ability. These gel-shift experiments used the different probes recognized by either or both PD and HD domains and show similar *in vitro* binding activity for all deletion mutants, including Pax7-291 that is deleted of the entire C-terminus (Figure 5b). The smallest C-deletion mutant, Pax7-465, is deleted of C-terminal sequences that are unique to Pax7 compared to Pax3 and that contain the OAR motif. Pax7-465 exhibits similar recruitment by ChIP-Seq as the reference Pax7-503 at pioneered sites (Figure 5c) leading to similar DNA accessibility measured in ATAC-Seq (Figure 5d) and deposition of similar bimodal distribution of H3K4me1 (Figure 5e). The OAR motif is thus not required for pioneer ability, although activity might be slightly decreased. Further C-terminal deletion to position 431 leads to loss of recruitment at ∼60% of Pioneer sites (Figure 5c), but not to Constitutive sites. Finally, almost complete deletion of the C-terminus to position 291 abrogates recruitment to Pioneer sites and reduces recruitment to Constitutive sites (Figure 5c) despite intact *in vitro* (EMSA) binding (Figure 5b).

**Figure 5.**
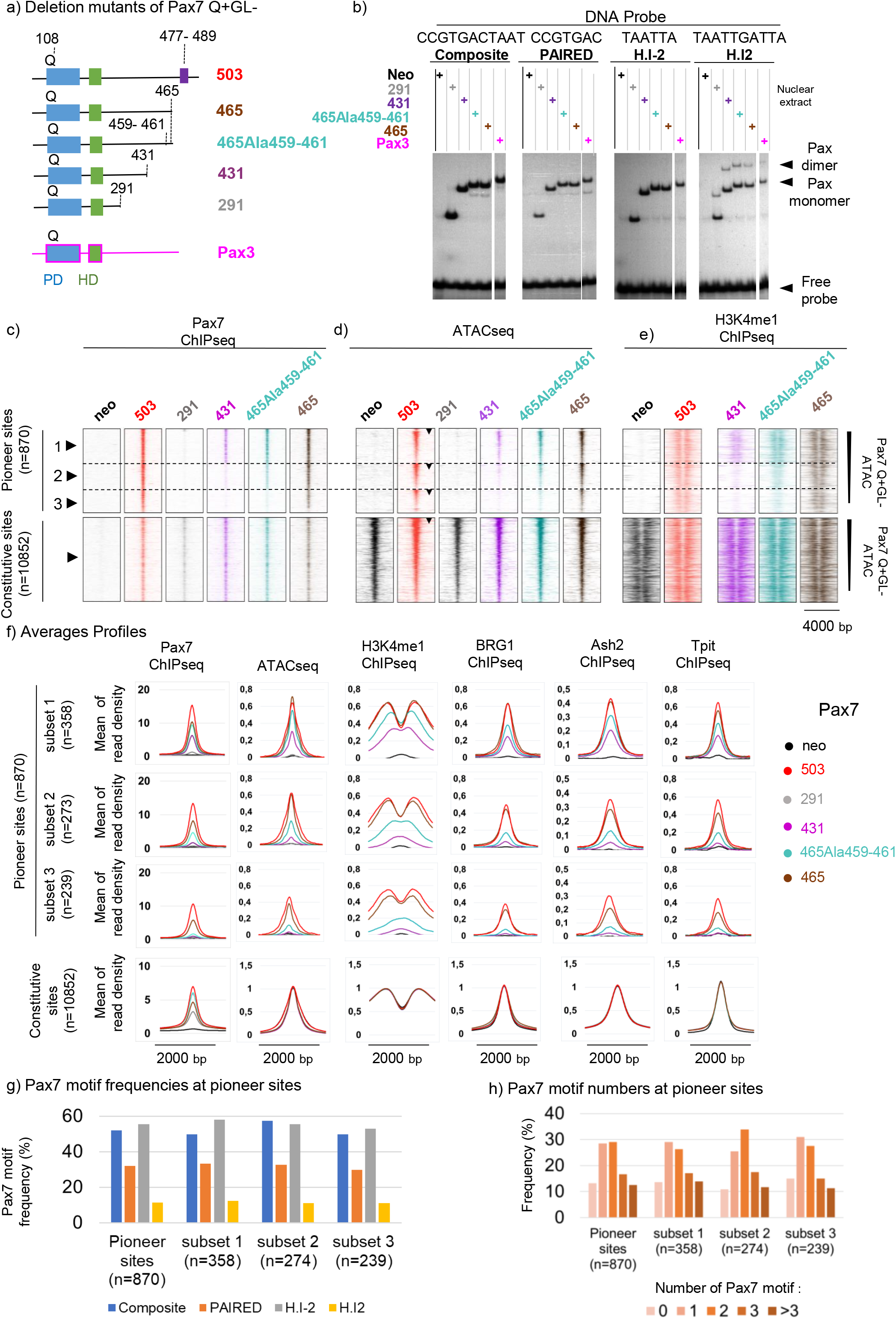
Pax7 amino acids 431-465 are necessary for pioneer activity. **a**) Schematic representation of C-terminal deletion mutants of Pax7 assessed for pioneer ability. **b**) In vitro DNA binding activity of Pax7 C-terminal deletion mutants assessed by EMSA using the indicated DNA probes. **c-e**) Chromatin opening ability of Pax7 C-terminal deletion mutants assessed by Pax7 ChIP-Seq (**c**), ATAC-Seq (**d**) and H3K4me1 ChIP-Seq (**e**). Subsets of Pioneer peaks that are impaired at different levels are indicated by dashed lines and numbered on the left. **f**) Average profiles for Pax7 ChIp-Seq, ATAC-Seq, H3K4me1, BGR1, Ash2 and Tpit ChIP-Seq for the subsets of Pioneer sites affected by C-terminal deletions. **g**,**h**) Frequencies (g) and total numbers (h) of Pax7 binding DNA motifs present at different subsets of Pioneer sites defined by Pax7 C-terminal deletions.

The critical interval between positions 431 and 465 is interesting because it is quite conserved compared to Pax3 and because it contains multiple putative sites of phosphorylation (Supplementary Figure 6a). To identify critical residues within this interval for Pax7 recruitment to pioneer sites, we performed a series of alanine replacement mutations throughout the interval and assessed those Pax7 mutants (Supplementary Figure 6b-d). Mutagenesis of either residues 459-461 or 463-464 to alanines had similar deleterious effects on recruitment to the *Pcsk2* pioneer site (Supplementary Figure 6c) but not to a transcriptionally activated site (*Pde2a*), and corresponding effects on *Pcsk2* mRNA levels (Supplementary Figure 6d). These mutations have marginal effects on a gene, *Lmcd1*, that is transcriptionally activated (ie not requiring pioneer action) by Pax7 (Supplementary Figure 6d). When assessed by ChIP-Seq, Pax7-465Ala^459-461^ shows a subset of sites (Figure 5c, subset 2) that has reduced but significant Pax7 binding in contrast to a subset 3 that shows no recruitment as for Pax7-431 (Figure 5c). As would be expected, the Pioneer sites that are no longer recognized by Pax7-431 or Pax7-465Ala^459-461^ (subset 3) show no ATAC signal (Figure 5d) and no H3K4me1 deposition (Figure 5e, f). In contrast, subset 2 of peaks that retain reduced Pax7 recruitment with Pax7-465lAla^459-461^ exhibits reduced ATAC-Seq signals and deposition of H3K4me1 with a single peak rather than bimodal distribution (Figure 5d, e). For Pax7-431, subset 2 shows no recruitment, ATAC or H3K4me1 signals, suggesting that the 431-465 interval contributes to chromatin interaction. Finally, subset 1 (peaks that have reduced Pax7 recruitment and ATAC-Seq signals for Pax7-431 and Pax7-465Ala^459-461^) shows a reduced single peak of H3K4me1 for Pax7-431 and a diminished bimodal signal for Pax7-465Ala^459-461^ (Figure 5c-f). The average profiles for these marks as well as for Tpit, Ash2 and Brg1 vary proportionately with the impact of deletions (Figure 5f). Thus, both Pax7-431 and Pax7-465 Ala^459-461^ behave similarly in the sense that for the subset of peaks where they retain weaker recruitment by ChIP-Seq, this is associated with reduced ATAC-Seq signals and a shift from bimodal to a single weaker peak of H3K4me1, typical of Primed enhancers. It is noteworthy that the three subsets of sites variably affected by Pax7 C-terminal deletions have similar frequencies of Pax7 binding motifs (Figure 5g) and total numbers of motifs (Figure 5h), suggesting that chromatin interaction may be the primary determinant for their recruitment.

This behavior is reminiscent of sites that are affected by the insertion of GL in the Pax7 Q+GL+ isoform (Figure 3). Comparison of the GL-sensitive subsets of Pioneer sites with those affected by the Pax7-465Ala^459-461^ showed that the former subset is overlapping with sites that show the loss of Pax7 recruitment or a shift from H3K4me1 bimodal to single peak with loss of ATAC signal following C-terminal deletion (Figure 6a-e). Hence, enhancers that exhibit impaired chromatin opening together with Primed rather than Active state because of GL+ insertion in the PD domain are also dependent on C-terminal sequences between 459-465 for chromatin remodelling. Overall, the Pax7-465Ala^459-461^ mutant shows impairment at a greater number of sites compared to the GL+ isoform but the latter are all included in the former group. The dependence of the same subset of enhancers on these two structural features of Pax7 for pioneer action may suggest that these enhancer’s contexts are similar. Further, these enhancer subsets exhibit parallel losses of Tpit recruitment (Figure 6f). In view of the importance of Tpit recruitment for complete enhancer opening (2), the reduced recruitment of Tpit at GL-sensitive enhancers can account for the observed blockade at the Primed state.

**Figure 6.**
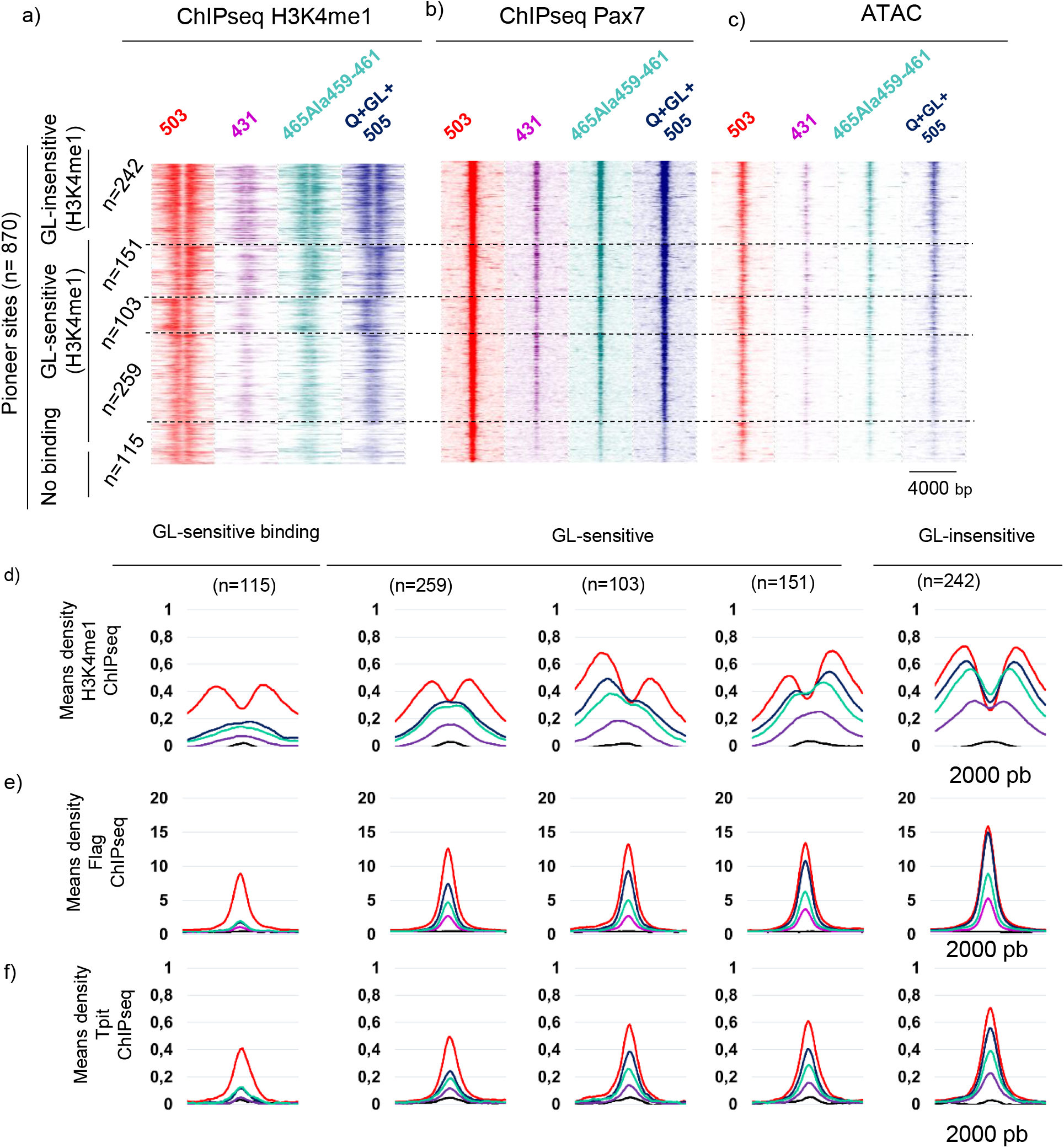
Pax7 paired domain GL+ insertion and amino acids 431-465 have similar impacts on melanotrope enhancer opening. **a-c**) Comparison of the deleterious effects of Pax7 C-terminal deletions with insertion of GL in the paired domain of isoform Q+GL+. The similarities are present in H3K4me1 (a) and Pax7 (b) ChIP-Seq, as well as in ATAC-Seq (c) profiles. Subsets are presented as defined by impact of GL+ isoform (Figure 3). **d-f**) Average profiles of indicated chromatin marks at GL-sensitive subsets of Pioneer peaks for Pax7 isoforms and critical C-terminal deletions as indicated by color-coding of Pax7 forms in (a).

## Discussion

We showed in the present work that the GL+ isoform of Pax7 produced by differential splicing of an extra two amino acids into the DNA binding PD domain loses significant pioneer ability without any effect on its transcription activation potential. The loss of pioneer ability is not complete as it affects a subset of its Pioneer sites in AtT-20 cells. The affected subsets include one (about 13% of Pioneer sites) that loose Pax7 recruitment in ChIP-Seq experiments and other subsets (three GL-sensitive subsets alltogether representing 59% of Pioneer sites) that fail to be completely opened and remain in the Primed state (Figure 3). Nonetheless, these subsets are significant as they lead to failure to activate most of the melanotrope transcriptome. The similar transcriptional activity of the various Pax7 isoforms (Figure 1e) suggests that they may be redundant for transcriptional maintenance. However, the isoform-specific pioneering ability implies that implementation of new enhancer repertoires by Pax7 during development may be linked to production of the appropriate isoform, the GL-isoform in the pituitary. While the different isoforms of Pax7 are expressed in adult tissues, there is no documentation of these differential splicing events at critical times during either pituitary, muscle or brain development.

Another differential splicing of the *Pax7* gene appears crucial for function: indeed, a splicing mutant resulting in loss of the OAR-containing isoform of human PAX7 is associated with a global neurodevelopmental delay (33). This mutation results in production of PAX7 isoforms that are similar to Pax7-465 terminating with coding sequence of the Pax7 exon 8. Since our analyses (Figure 5) indicate that Pax7-465 is sufficient for pioneer ability and function in pituitary cells, this suggests that some PAX7 functions in neural tissues might be uniquely dependent on the OAR domain.

Could relative affinity for genomic target sites be the limiting factor that accounts for loss of pioneering ability by the GL+ isoform and/or the C-terminal deletion mutants? A slightly lower apparent affinity of the GL+ isoform for target DNA in EMSA (Figure 2e) could reflect such dependence. However, some of the more severely affected Pax7 deletion mutants show decreased genomic recruitment by ChIP-Seq but no loss of *in vitro* DNA binding (EMSA) (Figure 5b). In contrast, the different subsets of Pioneer sites sensitive to either GL+ (Figure 3) or to C-terminal deletions (Figure 5) show overlapping distributions of Pax7 ChIP-seq recruitment strength and ATAC signals. So, these distributions argue against DNA binding strength as a primary determinant of pioneer ability (Figure 6a). Also, the apparent Pax7 ChIP recruitment signal intensity in itself is not well correlated with chromatin opening ability. A similar conclusion was reached by analysis of initial recruitments at different Pax7 pioneer sites where initial recruitment strength is inversely correlated with the levels of linker histone H1 (2). But it could be a combination of DNA and chromatin interaction strengths, together with ability to recruit the cooperating TF Tpit (as discussed below), that ultimately determine chromatin modification and opening. In this respect, it is noteworthy that Pax7-291 which is deleted of most C-terminal sequences with an intact DNA binding domain, binds DNA *in vitro* (Figure 5b) but fails to be recruited at Pioneer sites in chromatin (Figure 5b), highlighting the crucial role of the C-terminus for chromatin recruitment at closed sites but not at the already open Constitutive sites (Figure 5c). Within the Pax7 C-terminus, the progressive deletions suggest that sequences within both the aa291-431 and aa431-465 regions contribute to chromatin interaction at the closed Pioneer sites. Similarly, chromatin and linker histone H1 interaction of the pioneer PU.1 was mapped outside its DNA binding domain (16). In summary, despite some relationships between recruitment strength and pioneering ability, it is likely that other parameters are critical to explain the loss of pioneering ability.

It is noteworthy that the GL+ isoform or the deletion mutants that are impaired in pioneering ability exhibit subsets of targets that appear blocked in the primed state rather than being fully activated by the reference Pax7 (Figure 7). Blockade at the primed state is expected of a pioneering process going through the first step of chromatin opening but failing to undergo the second cell-cycle dependent step that involves nucleosome displacement and recruitment of the SWI/SNF complex (2). We also showed that this second step requires the cooperating nonpioneer factor Tpit in addition to cell replication (2). It is thus possible that pioneer ability of Pax7 mutants may result in the impaired ability of these mutants to recruit/interact with Tpit. The loss of Tpit recruitment at the GL-sensitive and Pax465Ala^459-461^ subsets of impaired pioneer sites is similar (Figure 6f) and hence this is a reasonable explanation. And indeed, the subset of sites that are only primed by Pax7 in AtT-20 cells does not recruit Tpit (2,8) and knockout of the *Tpit* gene results in failure of Pax7 pioneer capacity in the mouse pituitary (18). Further, Pax3 behaves very similarly to the GL+ Pax7 isoform in pituitary cells despite Pax3 being GL- and it also shows reduced Tpit recruitment (Figure 6f). This may be taken to suggest that the impact of GL insertion in Pax7 is at a tridimensional level, and this would be consistent with prior work suggesting that a unique conformation of Pax7 (that requires both intact paired and homeo domains) is involved in pioneering ability (13).

**Figure 7.**
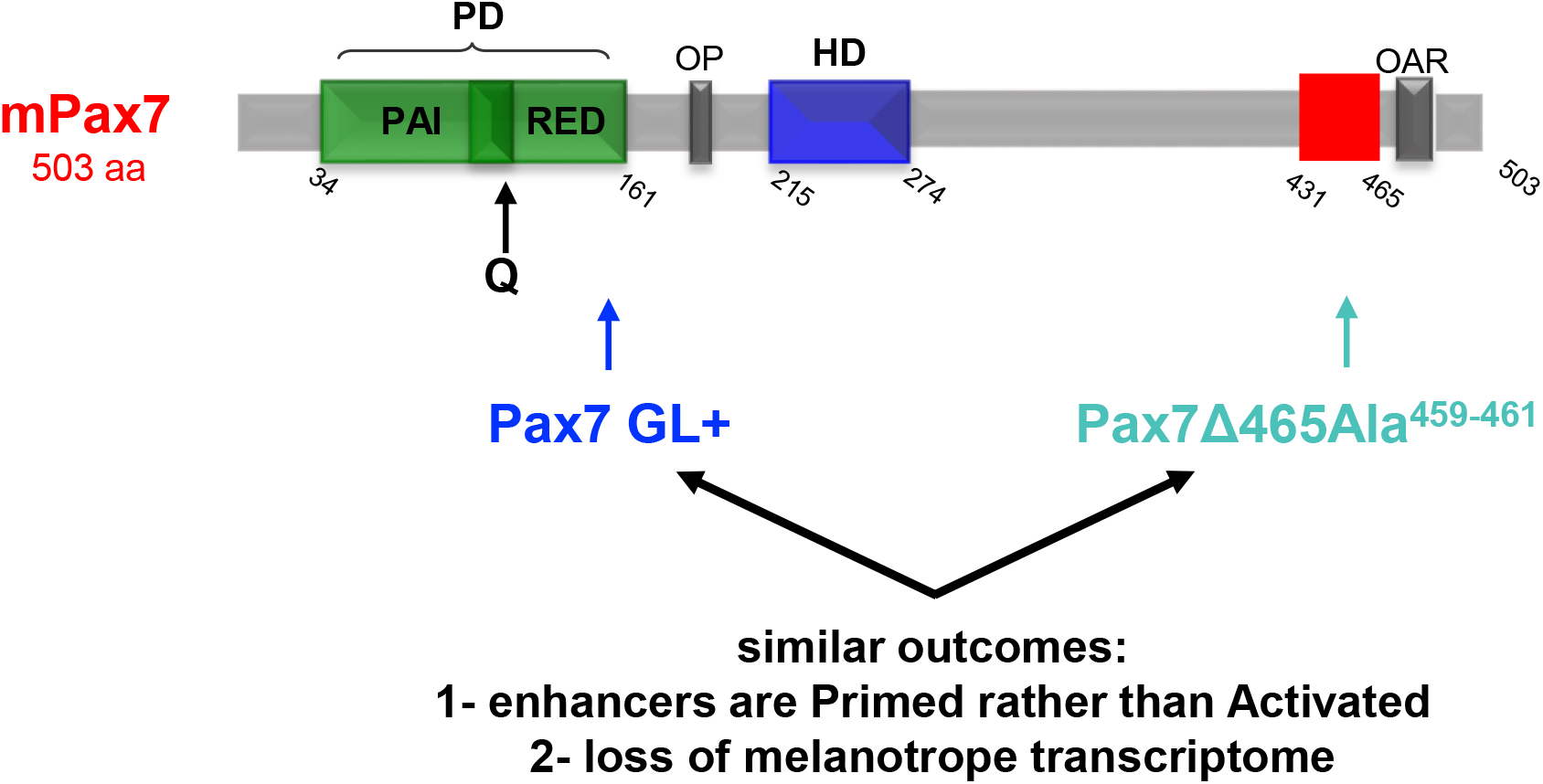
Pax7 features associated with impaired pioneer action and loss of melanotrope transcriptome. Schematic representation of mouse Pax7 alterations that prevent activation of large subsets of Pioneer sites and that are associated with activation of the melanotrope transcriptome. Both insertion of GL residues in a naturally-occurring isoform (GL+) and C-terminal deletion to position 465 together with alanine replacement of indicated conserved residues (Pax7del465Ala459-461) have very similar effects on pioneer activity.

Considered globally, the results present a picture of gradual loss of pioneering ability by the GL+ and deletion mutants that affect the same subsets of pioneer sites suggesting that it is the intrinsic properties of those sites that underlie their sensitivity. The nature of the critical properties that are involved remains to be better defined.

## Data Availability

Sequencing data are available under GSE87185 and GSE225231.

## Acknowledgements

We are grateful to Odile Neyret and Sarah Boissel, and their group at the IRCM Sequencing Core Facility for NGS analyses and to Valerie Magoon for her expert secretarial assistance.

