## Supplementary Figures for "Chromatin opening ability of pioneer factor Pax7 depends on unique isoform and C-terminal domain"

### Legends to Supplementary Figures

**Supplementary Figure 1. Characterization of Pax7 isoforms expressed in AtT-20 cells.**

**a**) Western Blot analysis of pools of AtT-20 cells expressing each Pax7 isoform (arrowhead) and showing similar levels of Pax7 protein compared to PCNA control (arrow).

**b**) PCA plot analysis of transcriptome data for RNA-Seq duplicates of AtT-20 cells expressing each Pax7 isoform and Pax3.

**c**) Volcano Plot representation of differentially expressed genes in AtT-20 cells expressing the reference Pax7 Q+GL- isoform compared to control Neo cells.

**d**) Volcano Plot representation of differentially expressed genes in AtT-20 cells expressing the Q+GL+ isoform compared to control Neo cells.

**e**) Transcriptional activity of Pax7 isoforms assessed by co-transfection with luciferase reporters driven by trimers of the indicated Pax7 binding sites inserted upstream of the minimal POMC promoter.

**Supplementary Figure 2. Characterization of ChIP-Seq replicates for Pax7 and its isoforms expressed in AtT-20 cells.**

**a**) Comparison of Pax7 ChIP-Seq signals for replicates of each Pax7 isoforms.

**b**) Comparisons of ChIP-Seq signals for each isoform compared to replica 1 of Pax7 reference isoform Q+GL-.

**c**) Means fold changes by site categories for each Pax7 isoform.

**Supplementary Figure 3. Isoform specific chromatin opening correlates with melanotrope transcriptome.**

**a-c**) Pioneer ability of the two pituitary-expressed Pax7 isoforms assessed by ATAC-Seq (**a**), Pax7 ChIP-Seq (**b**) and H3K4me1 ChIP-Seq (**c**). Heatmaps are shown for the subsets of Pioneer, Activated and Constitutive enhancer sites. The Pioneer subset is subdivided in two subsets of GL-sensitive (ATAC) and GL- insensitive (ATAC) based on the ATAC-Seq profiles.

**d**) Comparisons of Pax7 ChIP-Seq strength (reads density) for the two subsets of GL-sensitive and GL-insensitive pioneered sites.

**e**) Distribution of ATAC-Seq signals for Pax7 isoforms at the GL-sensitive and GL-insensitive Pioneer sites.

**f**) Frequencies of Pax7 binding motifs present in each subset of Pax7 Pioneer sites.

**g**) Total number of Pax7 binding motifs per subset of Pioneer sites.

**Supplementary Figure 4. Motif analysis for the GL related Pax7 Pioneer sites.** Searches were conducted for de novo and known DNA binding motifs as indicated.

**Supplementary Figure 5. Read density plots and correlations between marks at Pioneer sites**

**a**) Box plot comparisons of the Pax7 GL- and GL+ isoforms for recruitment of Pax7, Tpit, Ash2, Brg1 measured by ChIP-Seq (reads density) at the different subsets of GL-sensitive and GL-insensitive Pioneer sites.

**b**) Top -Correlation between the recruitment of the Pax7 isoform Q+GL+ and recruitment of the nonpioneer factor Tpit, Ash2 and Brg1 at Pioneer sites (reads density). Bottom- Correlation between the recruitment of Tpit and the remodeling factors Ash2 and Brg1 at Pioneer sites for the Pax7 isoform Q+GL+ (reads density).

**Supplementary Figure 6. Structure function investigation of Pax7.**

**a**) Sequence comparison of mouse Pax3 with Pax7. Paired and HD domains are highlighted in blue and green respectively, and the endpoints of C-terminal deletions indicated by red bar. The OAR domain is in purple. Yellow and light blue boxes indicate residues mutated to alanine.

**b**) Western Blot analyses of Pax7 C-terminal deletion-e mutants expressed in AtT-20 cells compared to PCNA control.

**c-d**) Assessment of different alanine mutations in the interval between aa431-465 of Pax7-465 for pioneer and transcriptional activity upon expression in AtT-20 cells. Genomic recruitment of the different mutants was assessed by ChIP-qPCR (**c**) and target gene expression by RT-qPCR (**d**) at a locus requiring enhancer opening by pioneer action (*Pcsk2*) and at loci where Pax7 acts only as a transcriptional activator (*Pde2a* and *Lmcd1*).

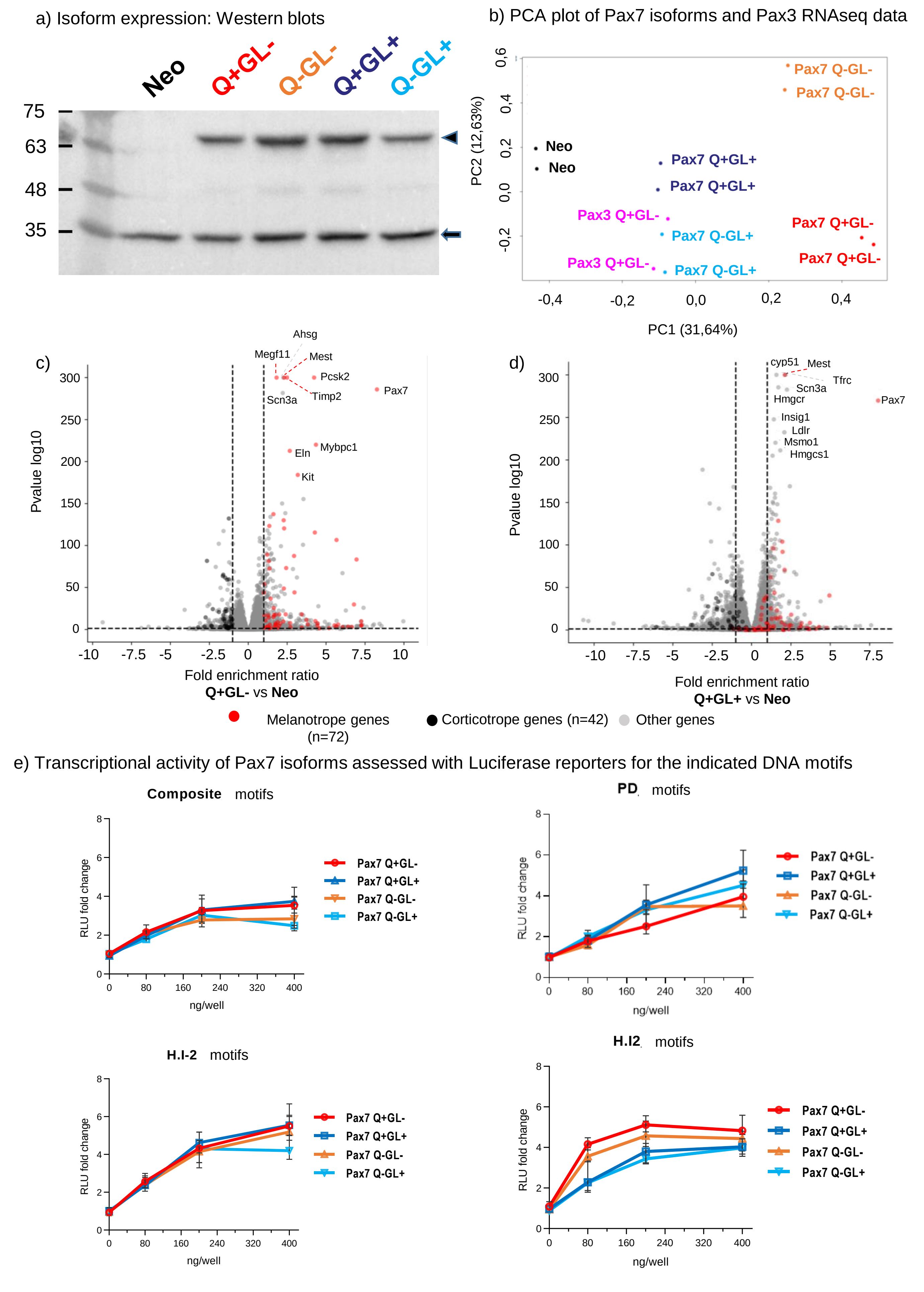


Supplementary Figure 1. RNAseq reproducibility and transcriptional activity

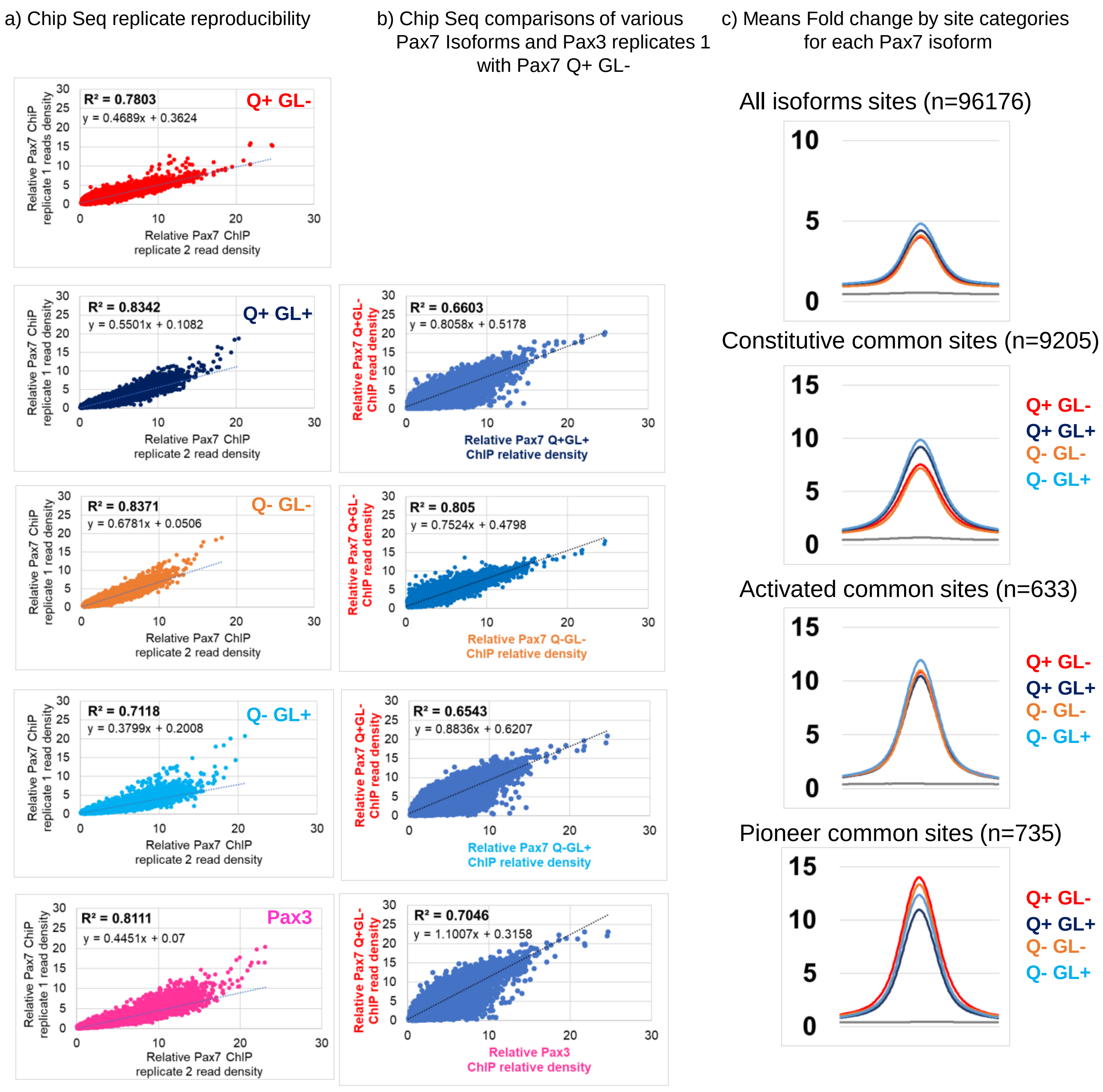


Supplementary Figure 2. ChIPseq reproducibility

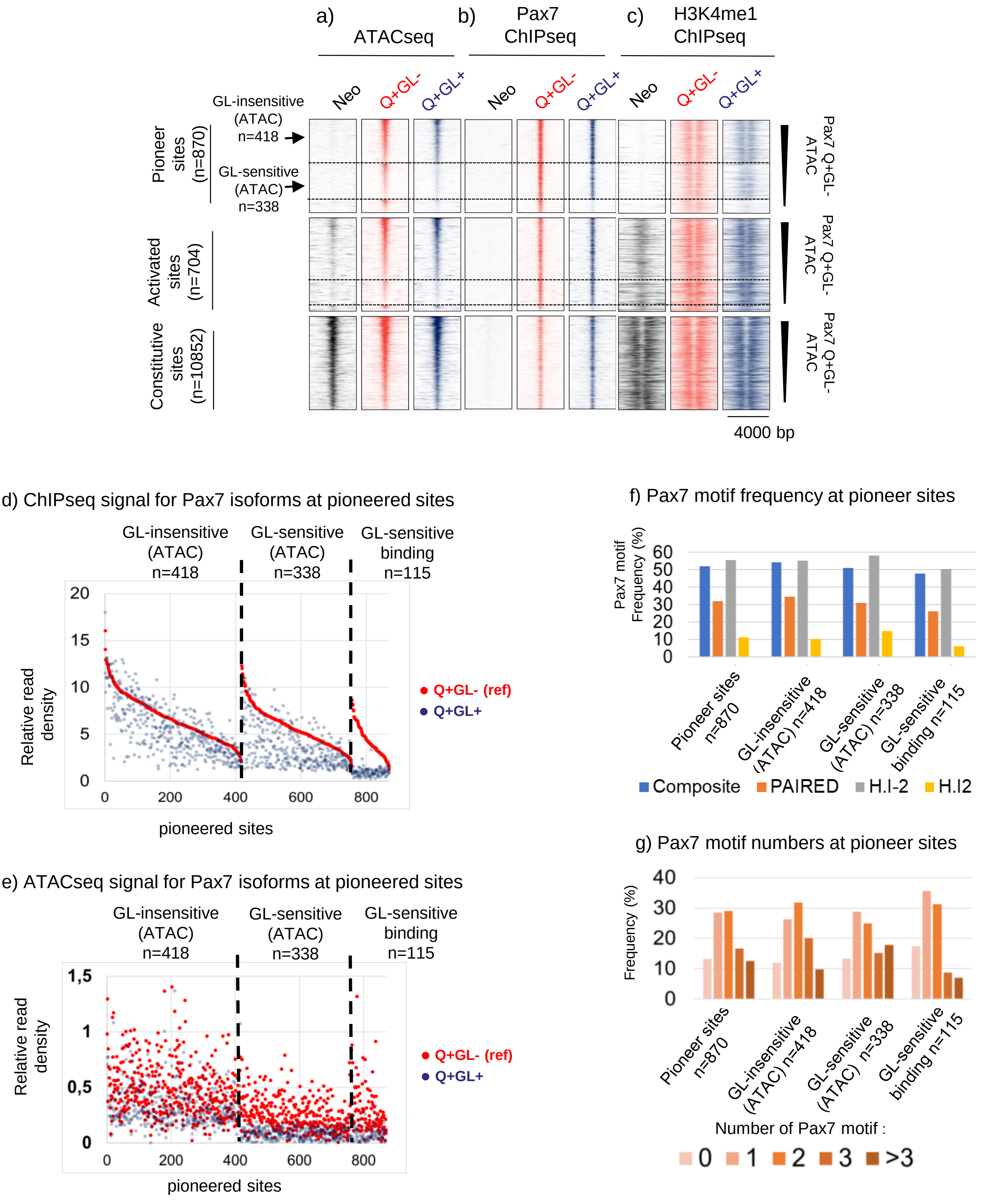


Supplementary Figure 3. Isoform-specific chromatin opening correlates with melanotrope transcriptome

### GL-insensitive (H3K4me1) n=242

#### De novo motifs

| Rank | Motif | P-value | % of Targets | % of Background | |
|---|---|---|---|---|---|
| 1 | 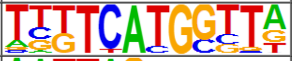 | 1e-44 | 44.21% | 9.41% | PAIRED related |
| 2 | 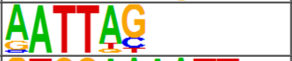 | 1e-28 | 78.51% | 43.23% | H.I2 |
| 3 | 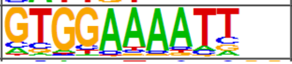 | 1e-25 | 34.71% | 9.63% | NFAT |

#### Known motifs

| Rank | Motif | Name | P-value | % of Targets Sequences with Motif | % of Background Sequences with Motif | |
|---|---|---|---|---|---|---|
| 1 | 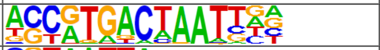 | PAX3:FKHR-fusion(Paired,Homeobox)/Rh4-PAX3:FKHR-ChIP-Seq(GSE19063)/Homer | 1e-27 | 20.66% | 2.80% | Composite |
| 2 | 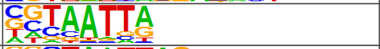 | DLX5(Homeobox)/BasalGanglia-Dlx5-ChIP-seq(GSE124936)/Homer | 1e-16 | 30.99% | 11.04% | H.I2 |
| 3 | 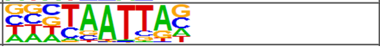 | DLX2(Homeobox)/BasalGanglia-Dlx2-ChIP-seq(GSE124936)/Homer | 1e-14 | 45.45% | 22.24% | H.I2 |

### GL-sensitive (H3K4me1) n=259

#### De novo motifs

| Rank | Motif | P-value | % of Targets | % of Background | |
|---|---|---|---|---|---|
| 1 | 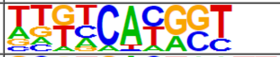 | 1e-56 | 55.60% | 13.47% | PAIRED related |
| 2 | 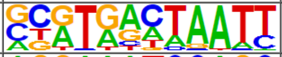 | 1e-39 | 32.05% | 5.43% | Composite |
| 3 | 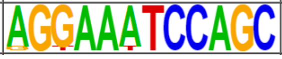 | 1e-29 | 4.25% | 0.01% | RBM6 |

#### Known motifs

| Rank | Motif | Name | P-value | % of Targets Sequences with Motif | % of Background Sequences with Motif | |
|---|---|---|---|---|---|---|
| 1 | 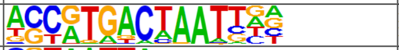 | PAX3:FKHR-fusion(Paired,Homeobox)/Rh4-PAX3:FKHR-ChIP-Seq(GSE19063)/Homer | 1e-37 | 24.32% | 2.93% | Composite |
| 2 | 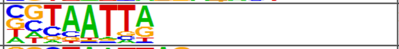 | DLX5(Homeobox)/BasalGanglia-Dlx5-ChIP-seq(GSE124936)/Homer | 1e-22 | 33.59% | 10.73% | H.I2 |
| 3 | 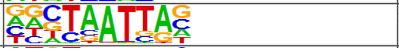 | En1(Homeobox)/SUM149-EN1-ChIP-Seq(GSE120957)/Homer | 1e-22 | 54.44% | 25.65% | H.I2 |

### GL-sensitive binding n=115

#### De novo motifs

| Rank | Motif | P-value | % of Targets | % of Background | |
|---|---|---|---|---|---|
| 1 | 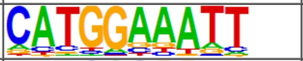 | 1e-18 | 49.57% | 14.39% | NFAT |
| 2 | 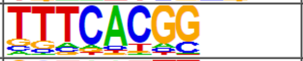 | 1e-15 | 61.74% | 25.48% | PAIRED related |
| 3 | 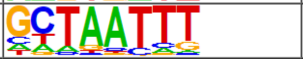 | 1e-12 | 55.65% | 24.04% | H.I2 |

#### Known motifs

| Rank | Motif | Name | P-value | % of Targets Sequences with Motif | % of Background Sequences with Motif | |
|---|---|---|---|---|---|---|
| 1 | 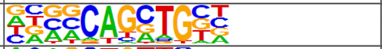 | Ascl1(bHLH)/NeuralTubes-Ascl1-ChIP-Seq(GSE55840)/Homer | 1e-14 | 46.96% | 15.92% | BHLH |
| 2 | 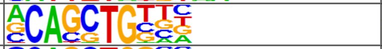 | Ptf1a(bHLH)/Panc1-Ptf1a-ChIP-Seq(GSE47459)/Homer | 1e-13 | 63.48% | 29.36% | BHLH |
| 3 | 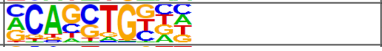 | HEB(bHLH)/mES-Heb-ChIP-Seq(GSE53233)/Homer | 1e-12 | 48.70% | 18.85% | BHLH |

Supplementary Figure 4. Motif analysis for the GL related sites

a) Read density plots of activating marks at pioneer enhancers

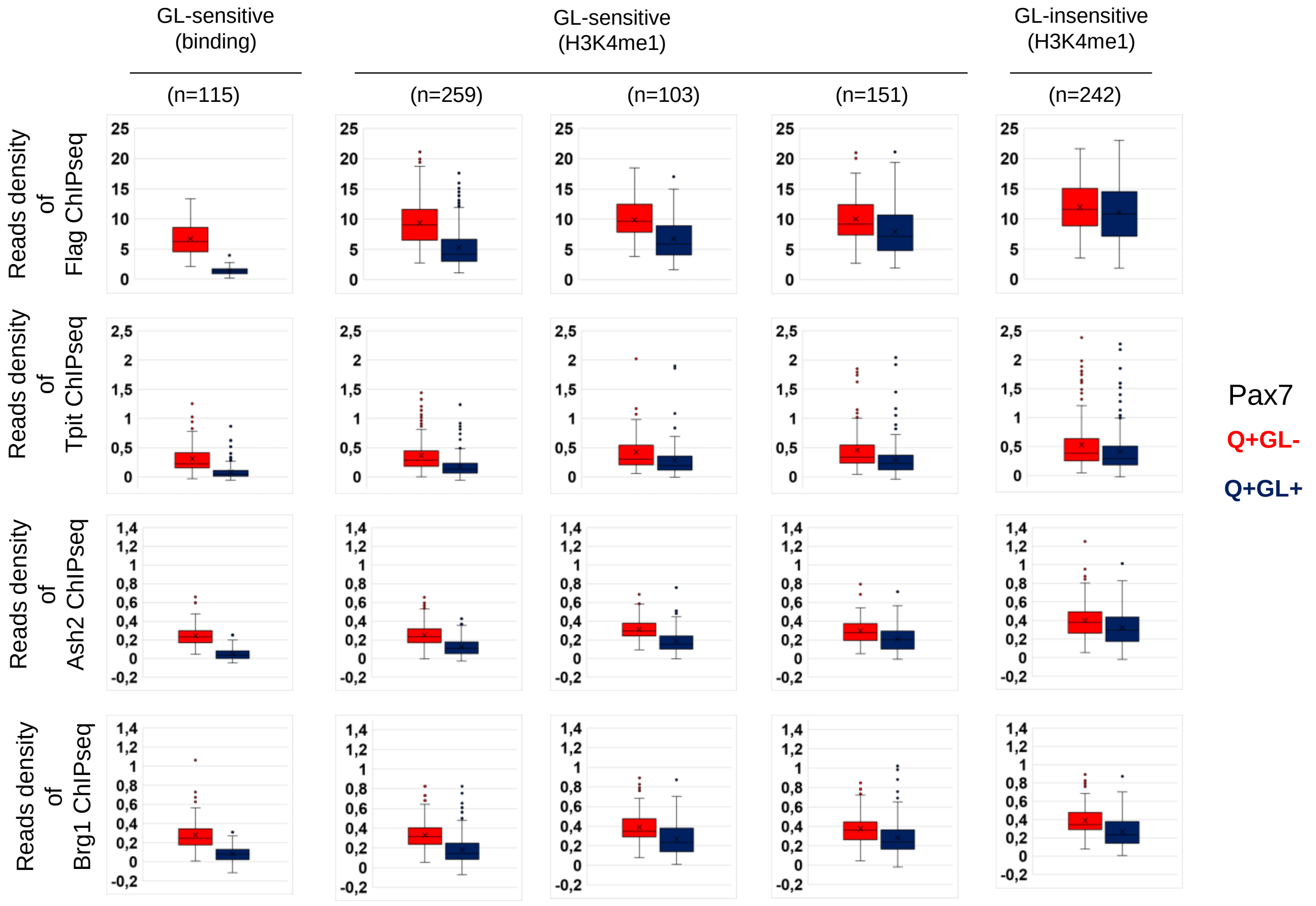


b) Correlation between activating marks and transcription factor recruitments

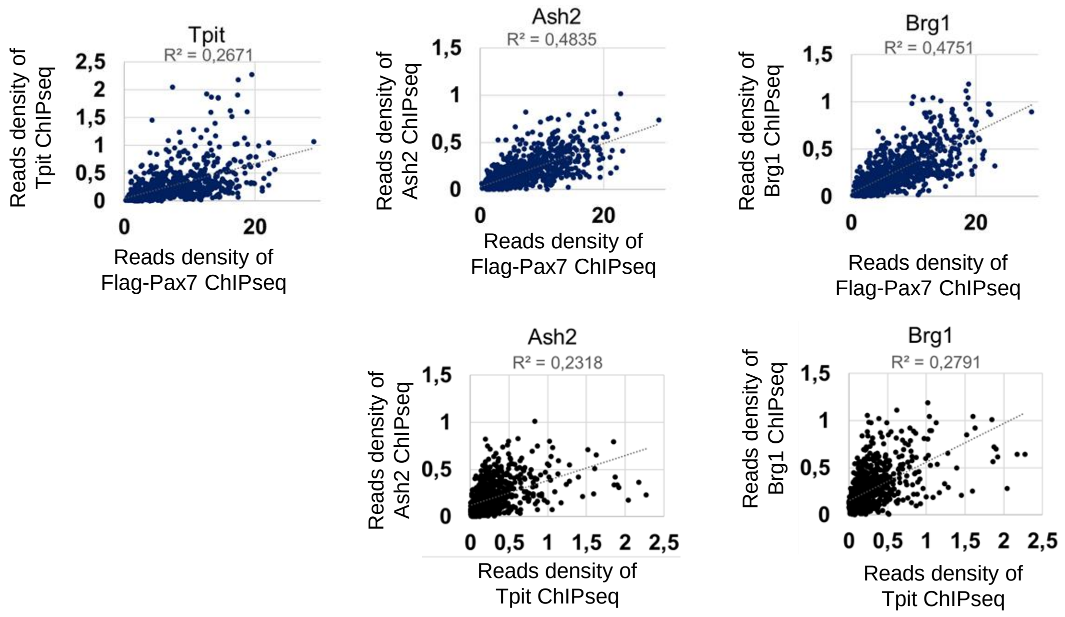


Supplementary Figure 5. Read density plots and correlations between marks at pioneer sites

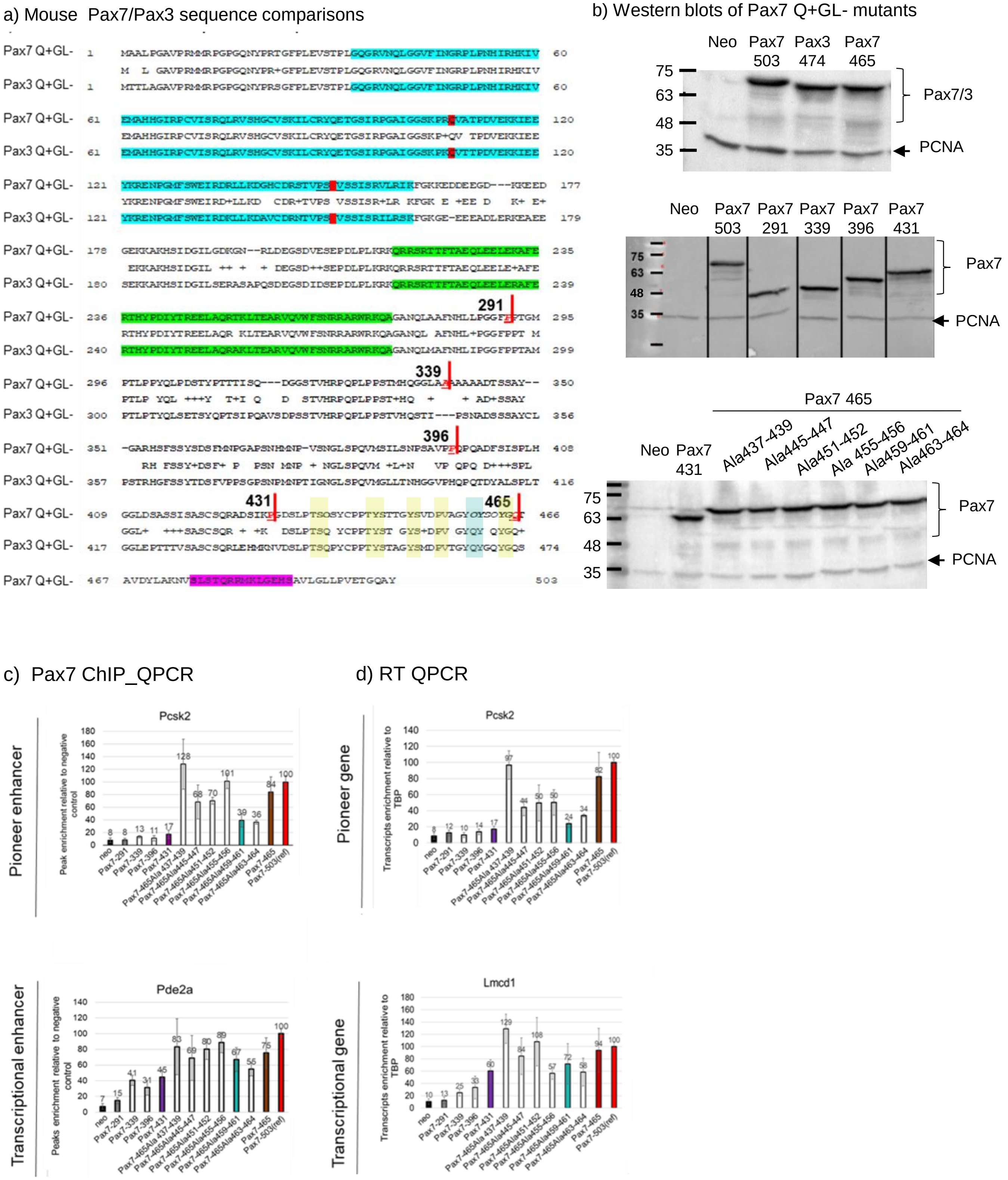


Supplementary Figure 6. Structure function study of Pax7
